# Quality of Life in Mothers of Children with Microcephaly in the Northeast of Brazil

**DOI:** 10.1101/663245

**Authors:** Caren Cristina Freitas Fernandes, Caíque Jordan Nunes Ribeiro, Anne Caroline Santos Melo de Araújo Cândido, Mariangela da Silva Nunes, Daniele Vieira Dantas, José Antonio Barreto Alves, Alanna Gleice Carvalho Fontes Lima, Heloísa Salvador dos Santos Pereira, Jonas Santana Pinto, Maria do Carmo Oliveira Ribeiro

## Abstract

**Objective:** To evaluate the quality of life of mothers of children with microcephaly compared to the quality of life of mothers with children of the same age but with normal neuropsychomotor development (NPM).

**Method:** This study was a cross-sectional, comparative, analytical study held in a public reference service. Seventy-eight (78) mothers with children between birth and two years old, with and without changes in their NMP, were interviewed. The abbreviated Questionnaire for the Evaluation of Quality of Life (WHOQOL-bref) and a sociodemographic evaluation questionnaire, developed by the author, were used. The data were analyzed descriptively, evaluating the association between variables and correlation tests.

**Results:** Mothers of children with microcephaly showed lower scores for various quality of life domains, however there was only a statistically significant difference for the environmental domain (48.40 for the group of mothers of children with microcephaly vs. 57.13 for the group of mothers with children with normal NPM, P<0.02). It should be noted that there were also significant negative correlations between the majority obstetric variables, maternal age and quality of life scores. There was no significant association between the child’s age and such scores.

**Conclusion:** Children with neuropsychomotor variations have not influenced their mother’s quality of life, rather, the mother’s quality of life is affected predominantly by housing conditions and financial resources.

**NON-TECHNICAL AUTHOR SUMMARY:** This study was a cross-sectional, comparative, analytical study held in a public reference service. Seventy-eight (78) mothers with children between birth and two years old, with and without changes in their NMP, were interviewed. The abbreviated Questionnaire for the Evaluation of Quality of Life (WHOQOL-bref) and a sociodemographic evaluation questionnaire, developed by the author, were used. The data were analyzed descriptively, evaluating the association between variables and correlation tests. Mothers of children with microcephaly showed lower scores for various quality of life domains, however there was only a statistically significant difference for the environmental domain (48.40 for the group of mothers of children with microcephaly vs. 57.13 for the group of mothers with children with normal NPM, P<0.02). It should be noted that there were also significant negative correlations between the majority obstetric variables, maternal age and quality of life scores. There was no significant association between the child’s age and such scores. Children with neuropsychomotor variations have not influenced their mother’s quality of life, rather, the mother’s quality of life is affected predominantly by housing conditions and financial resources.

## 1 INTRODUCTION

Microcephaly has been a worldwide concern since an increased incidence of the disease began in 2015, attracting the attention of several entities and the scientific community. However, few studies have evaluated the psychosocial aspects of the mothers of children affected by this condition.

Microcephaly is a congenital malformation in which the head and the brain of the child develop improperly, due to changes in morphology, physiology or metabolism of cells, tissues and/or organs. The etiology is related to morphological and biochemical abnormalities during pregnancy. Therefore, it can be diagnosed in the pre- or postnatal period. Recently, the World Health Organization (WHO) has updated the parameters for the diagnosis of microcephaly to include the following: cephalic perimeter equal to or less than 31.9 centimeters for boys, and equal to or less than 31.5 centimeters for girls [1, 2].

The cases of babies born with this malformation have been increasing since April 2015 in Brazil, and research indicates a possible association with Zika virus (ZIKV) [3]. In October 2015, an unexpected increase in newborn babies with microcephaly was noticed, especially in the northeast of Brazil. These numbers advanced so fast that by September 2016, according to the Ministry of Health (MH), 6,614 suspected cases of microcephaly had been registered, in which 1,989 were confirmed and 4,625 were discarded. During this time, there was also laboratory confirmation of the autochthonous circulation of the ZIKV in 22 federative units [4, 5]. In October 2016, 124 cases were confirmed in Sergipe [5].

Based on the previous statements, it is necessary to pay attention to the children’s mother’s health, since they are their main caretakers. Having a child is one of the most important events in a woman’s life. For this reason, when the mother is informed about the existence of a congenital malformation, her concerns for the future can increase considerably. These concerns accompany the mother throughout her life, with greater or lesser intensity, since such a disability is invariably associated with suffering, discomfort, and doubts, and requires a long-term investment, impacting the mother’s quality of life [6].

Quality of life (QL) is defined by the WHO Quality of Life Group as “the individual’s perception of their position in life in the context of the culture and value system in which they live and in relation to their goals, expectations, standards and concerns” [7]. A recent preliminary study performed in a neonatal intensive care unit (NICU) in Sergipe revealed that mothers of children with ZIKV-related microcephaly had high levels of anxiety and low scores for the psychological domain in the first 24 hours after birth when compared to a control group formed by mothers of healthy newborns [8].

This is the first study to investigate the QL of mothers of infants with microcephaly in our country. Our main objective was to evaluate the quality of life of mothers of children with microcephaly and compared to the quality of life of mothers with children of the same age but with normal neuropsychomotor development (NPM).

## 2 MATERIALS AND METHODS

This is a cross-sectional, comparative, analytical and quantitative study performed in a public reference outpatient service for follow-up cases of malformation in Aracaju, Sergipe, Northeast Brazil, from October 2016 to January 2017.

The sample was non-probabilistic and composed of 39 mothers of children with microcephaly (G1 group) and 39 mothers of children with eating disorders (G2 group) matched by age.

The G2 group was chosen due to the location of care and because the follow-up was the same for children with microcephaly, and due to the peculiarities of these children’s health problems. In eating disorders, only the gastrointestinal system is reached without compromising neuropsychomotor development. Food intolerance or eating disorders are any abnormal response of the body to the ingestion of a food, which may or may not trigger an immune response.

The inclusion criteria used were mothers of children between birth and to two years of age who were evaluated in our institution. Exclusion criteria were mothers who were psychologically unable to provide reliable data and mothers of children with normal development that had any associated pathology with a neuropsychomotor impairment.

The mothers were invited to join the research during breaks in their children’s specialized monitoring appointments, on different days and times. Sociodemographic and clinical data of these women and their children were collected through a standardized form. The QL was evaluated through The Questionnaire for the Evaluation of Quality of Life (WHOQOL-bref). The forms were filled out by the researchers in order to minimize gaps in understanding of information, due to the complexity of the investigated phenomenon and the participants’ varied schooling background.

WHOQOL-bref is a questionnaire of 26 questions, where the first two questions are about general QL and the other 24 represent each of the 24 facets that make up the original tool, each facet being evaluated by only one question. It is divided into 4 domains for QL analysis: Physical, Psychological, Social Relations, and Environment [9].

The participants were interviewed individually by the researchers without any interference in their responses, allowing time for each subject to respond. They were also informed about the procedures adopted during the research, including the risks and benefits. The study was approved by the Research Ethics Committee of the Federal University of Sergipe and followed the recommendations of the Declaration of Helsinki.

The data were analyzed descriptively and the Kolmogorov-Smirnov test was used to check the distribution symmetry. The parametric quantitative variables were represented by mean ± standard error of the mean and non-parametric variables were expressed as medians and percentiles 25 and 75 (p25, p75). Categorical variables were expressed in absolute and relative frequencies.

The association between categorical variables was assessed using the Chi-square test and Fisher’s Exact test and the association between quantitative variables was assessed using the Spearman and Pearson correlation tests. The Mann-Whitney U test and the student’s independent T test were used to verify the differences between the groups. Values of p <0.05 were considered statistically significant.

### Ethics Statement

The study was approved by the Research Ethics Committee of the Federal University of Sergipe and followed the recommendations of the Declaration of Helsinki.

## 3 RESULTS

Seventy-eight (78) women were part of this study (39 in each group). They were predominantly young adults (27.99±0.75 years old), with six to eleven years of schooling (56.4%), predominantly non-white não brancas (76.9%), with permanent partners (85.9%), from the state’s countryside (57.7%), with their own residence (70.5%). Their profession was often not mentioned (61.5%), and they often had a family income between one and two minimum wages (73.1%) and the median number of dependents was four people. Almost half of the participants did not receive government assistance (51.3%) (Table 1).

**Table 1.**
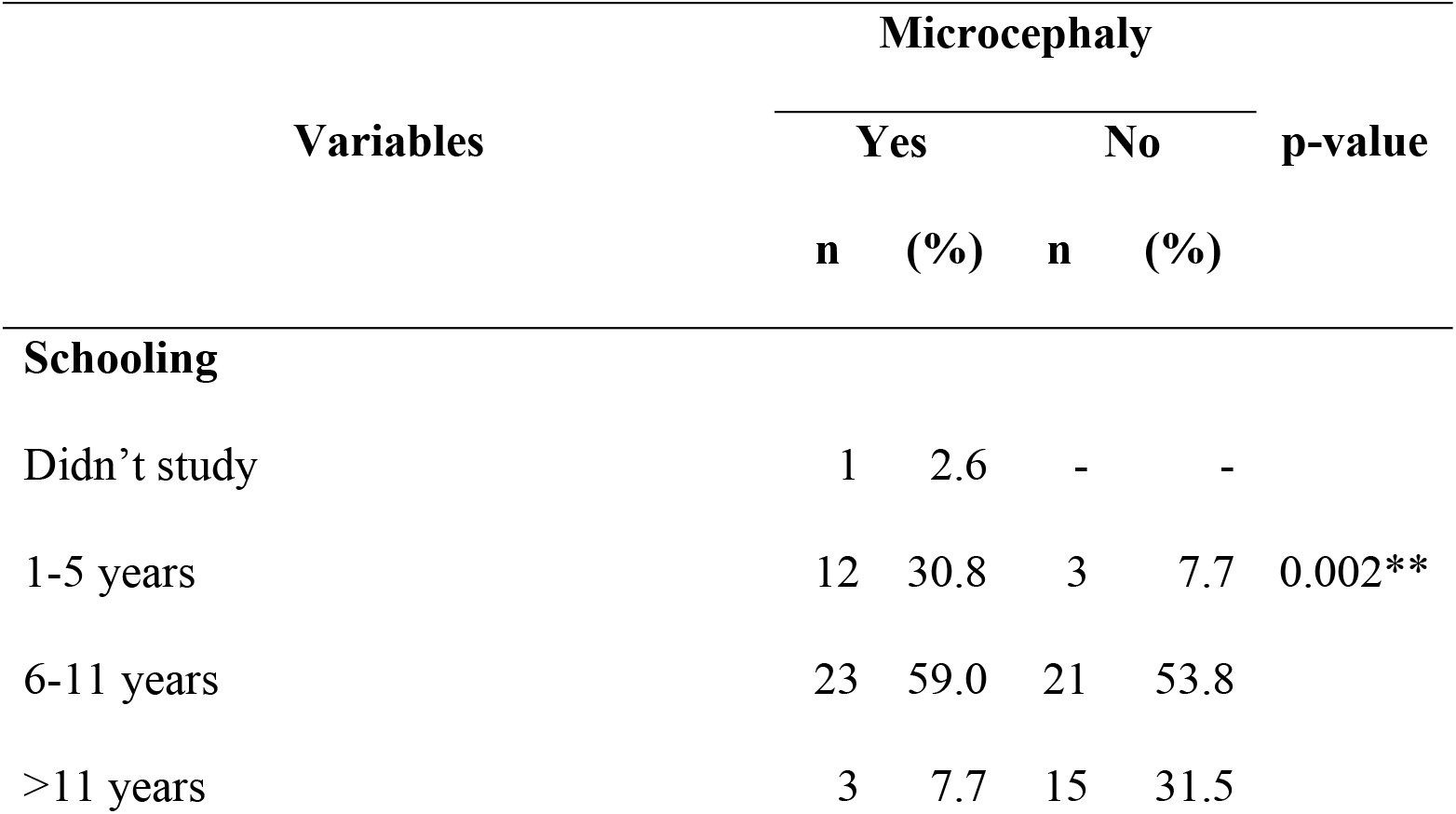
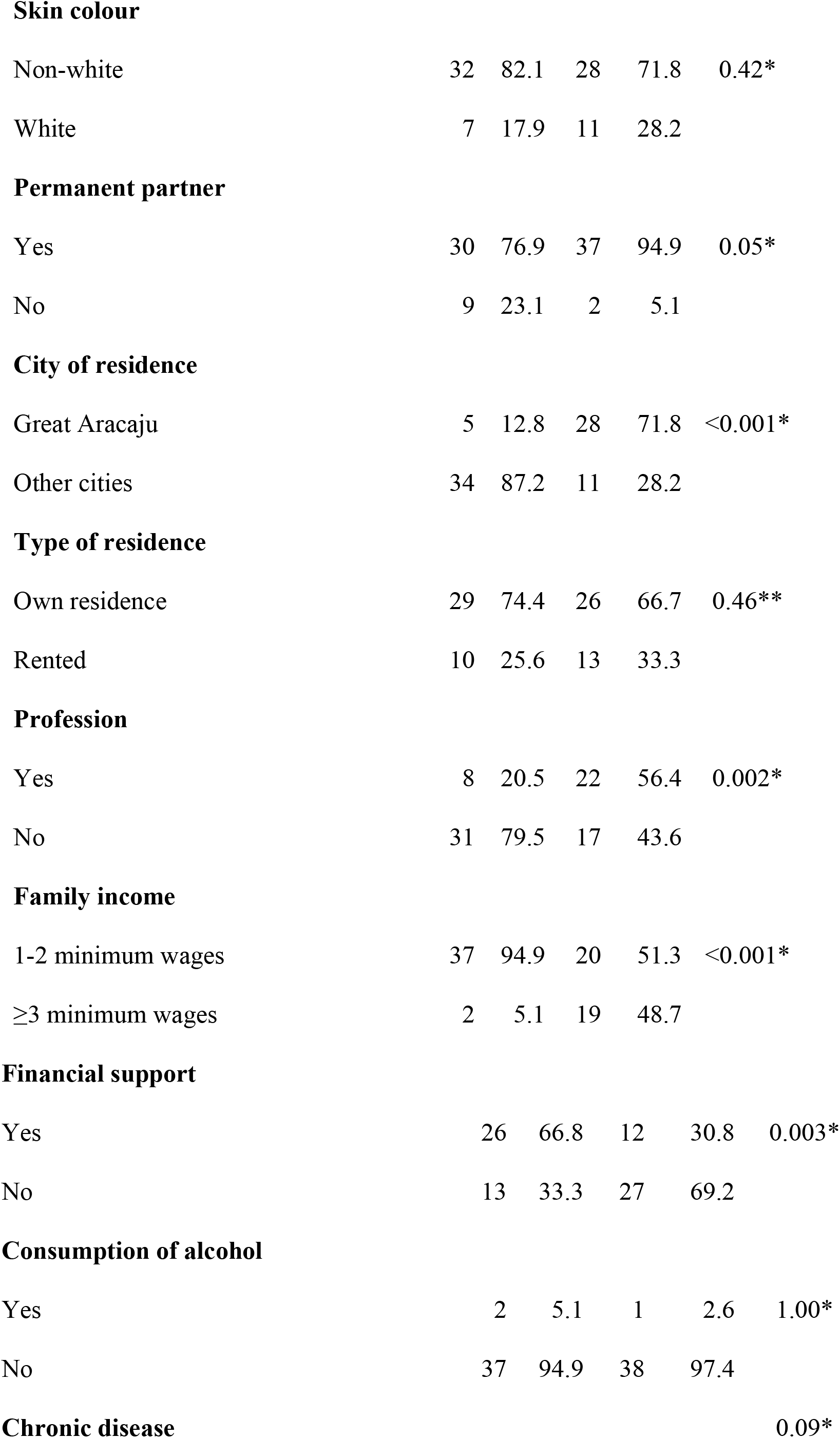
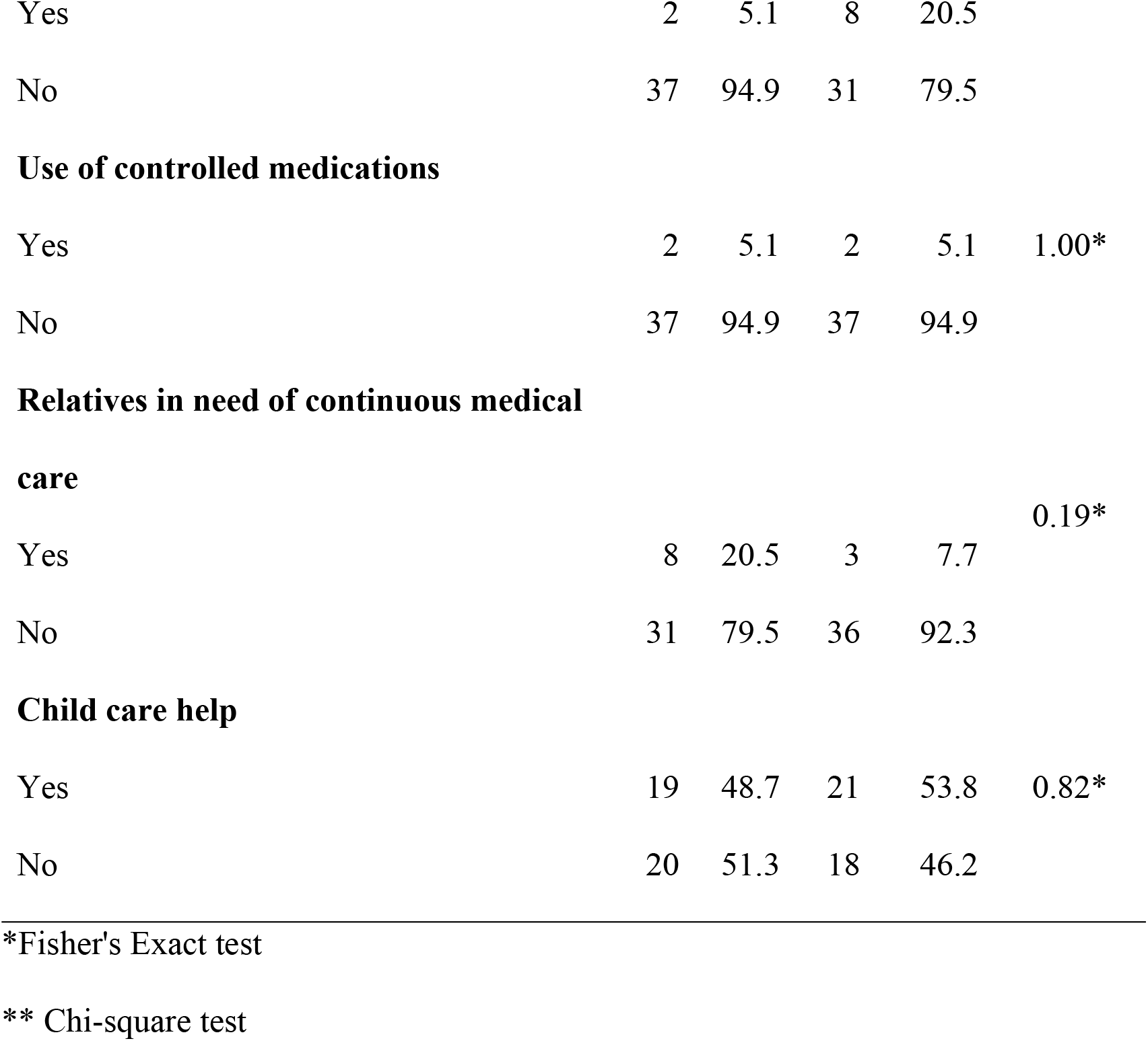
Sociodemographic characteristics. 2016-2017.

Most of the participants stated that they did not consume alcohol (96.2%), did not have any comorbidities (86.1%), did not use controlled medications (94.9%) and have no other relatives dependent on their care (85.9%). Only half had support from others for childcare (Table 1).

Participants had a median of two pregnancies, two births, two living children and no previous abortions. The sample was homogeneous regarding the majority of sociodemographic characteristics. However, significant differences were found between the groups when comparing the following characteristics: family income, municipality of residence, schooling, profession, financial aid, and child’s age and mother’s age. (Table 2).

**Table 2.** Social and obstetric characteristics. 2016-2017.

| Variables | Microcephaly | p-value |
| --- | --- | --- |

|  | Yes | No |  |
| --- | --- | --- | --- |
| **Age** | 26,23±1,09<sup>†</sup> | 29.74±0.96<sup>†</sup> | 0.02* |
| **Income dependents** | 4(3;5)<sup>††</sup> | 4(3;4)<sup>††</sup> | 0.13** |
| **Number of children** | 2(1;3)<sup>††</sup> | 2(1;2)<sup>††</sup> | 0.17** |
| **Number of pregnancies** | 2(1;4)<sup>††</sup> | 2(1;3)<sup>††</sup> | 0.42** |
| **Number of abortions** | 0(0;1)<sup>††</sup> | 0(0;1)<sup>††</sup> | 0.88** |
| **Number of births** | 2(1;3)<sup>††</sup> | 2(1;2)<sup>††</sup> | 0.32** |
| **Age of the child** | 10.46±0.35<sup>†</sup> | 8.10±0.96<sup>†</sup> | 0.01* |

<sup>†</sup>Media±EPM
<sup>††</sup>Medians (p25;p75)
\*Student t test(independent)
\*\*Mann-Whitney U. test

Although G1 mothers had lower QOL scores, there was a statistically significant difference only for the environmental domain (Table 3).

**Table 3.** Quality of life per group of caretakers. 2016-2017.

| Variables | Microcephaly |  | p-value* |
| --- | --- | --- | --- |
|  | Media±EPM |  |  |
|  | Sim | Não |  |
| Physical domain | 58.15±2.81 | 60.35±2.48 | 0.56 |
| Psychological domain | 61.75±2.74 | 62.29±2.51 | 0.89 |
| Social domain | 64.96±3.69 | 66.67±2.75 | 0.71 |
| Environmental domain | 48.40±2.59 | 57.13±2.69 | 0.02 |
| Total score | 58.31±2.47 | 61.61±1.99 | 0.30 |

\* Student t test(independent).

Significant negative correlations were observed between the majority of obstetric variables, maternal age and QOL scores. The age of the child was not significantly associated with such scores (Table 4).

**Table 4.** Correlation between the characteristics of the caregiver and the QL. 2016-2017.

| Variables | Physical | Psychological | Social | Environmental | Total score |
| --- | --- | --- | --- | --- | --- |
|  | domain | domain | domain | domain |  |
| Coeficiente de correlação (p-valor) |  |  |  |  |  |
| **Number of children** | -0.50<br>(0.001)* | -0.38 (0.017)* | -0.40<br>(0.011)* | -0.53 (0.001)* | -0.52<br>(0.001)* |
| **Number of pregnancies** | -0.43<br>(0.007)* | -0.39 (0.014)* | -<br>0.34(0.034)* | -0.54 (<0.001)* | -0.49<br>(0.001)* |
| **Number of abortions** | -0.07<br>(0.672)* | -0.03 (0.831)* | 0.19<br>(0.257)* | -0.07 (0.657)* | 0.01 (0.954)* |
| **Number of births** | -0.49<br>(0.001)* | -0.40 (0.012)* | -0.42<br>(0.008)* | -0.56 (<0.001)* | -0.55<br>(<0.001)* |
| **Woman's age** | -0.23<br>(0.163)** | -0.22<br>(0.173)** | -0.25<br>(0.123)** | -0.42 (0.008)** | -0.33<br>(0.041)** |
| **Child's age** | -0.02<br>(0.903)** | -0.16 (0.332)<br>** | -0.17<br>(0.292)** | 0.06 (0.730)** | -0.1<br>(0.548)** |

\* Spearman Correlation
\*\* Pearson's Correlation

## 4 DISCUSSION

Microcephaly currently represents a serious public health problem, impacting the QL of the mothers of these children due to comorbidities associated with the disease and the overload of care. The study evaluated the QL of 78 women, 39 mothers of children with microcephaly and 39 mothers of children with normal NPM, who were evaluated in the outpatient clinic of a public university hospital in the capital of Sergipe.

Despite the peculiarities of neuropsychomotor development of children with microcephaly, their mothers’ overall QL scores were not significantly different from the mothers of children with normal neuropsychomotor development.

Our study evaluated the QL of the mothers of children with microcephaly due to the increasing number of cases in Brazil and in other countries, and the impact that the disease causes on the life of these mothers. Previous studies have evaluated the QL of parents or caretakers of children with developmental disabilities or disorders such as autism, Down’s syndrome and cerebral palsy [6,10,11], but no studies on the QL of the mothers of children with microcephaly were found.

The QL goes beyond the absence of disease and overcoming the difficulties encountered in daily life. It can be influenced by multiple factors such as safety, transportation, education and housing [12,13,14]. The results showed that the QL of the mothers of children with microcephaly transcended the absence of disease, being influenced mainly by housing conditions and absence of transportation, among other factors.

When these services are lacking in quality, they can have emotional consequences that reduce QL. This research has showed that transport is very important for the mothers of children with microcephaly, since they live far from where the treatment and follow-up of their children’s illness is performed.

Most of the participants live in the state countryside, especially those from group G1. The environment in this study, is a reference for the medical assistance of children with microcephaly who live in the state countryside.

Housing conditions are important social determinants of health. Normally, there is a better infrastructure in the capital city, in contrast to the countryside. From the analysis of the diagnosed cases from ZIKV in the northeast of Brazil, it can be inferred that the incidence of this viral infection is closely related to carrier proliferation and, consequently, to the health conditions of the area [15, 16]. To prevent microcephaly, it is essential to fight the mosquitos that transmit ZIKV, which is related to the disease and malformation.

The results showed that, due to the majority of mothers of children with microcephaly living in the countryside, and the housing being so precarious, their QL has also been influenced by the environment they are living in.

A small percentage of the interviewed mothers from the G1 group had a higher education level, and the majority did not declare any professional occupation, as opposed to the results found in group G2. This finding may be related to a greater need for childcare, due to the limitations associated with the disease.

Studies have confirmed that, in addition to microcephaly, children showed hearing and visual damage and neuropsychomotor deficits of varying levels, which may lead to cerebral palsy [10, 18]. The results of our study showed that, because they are the main caretakers, these women end up giving up work and academic activities. Similar results were found in another study of mothers of children with disabilities, who despite being able to be in the labor market as young adults, take full responsibility for all childcare [19].

The results showed that the mothers of children with microcephaly gave up living their own lives, focusing only on the child and household care. These changes may influence the social life of these mothers and their family income.

Few participants reported receiving childcare assistance. The involvement of someone, especially the father, in care was very small, which is similar to the results reported in another study that addresses family care, including parental involvement, in children with disabilities [19].

Although most of the participants had a permanent partner, the responsibility for childcare rested entirely on the mother, especially for vital needs such as food, stimulation, understanding, affection and care. When childcare needs are fulfilled only by the mother, she might be overburdened, affecting her QL.

On the other hand, some of the mothers stated that they had family members in the household who also needed continued medical care. Previous studies also showed that the “psychological well-being” of caretakers is affected when they are overburdened [10].

The presence of a third person whom the woman was also taking care of, can directly influence her daily burden. Being in charge of the care of more than one family member can lead to greater physical and emotional exhaustion. The results show that due to the child’s comorbidities, the mother may suffer from being overburdened. Additionally, the presence of another family member who requires care may cause more stress and burden on the mother’s life, influencing her psychological condition. Due to the mother’s dedication to the needs of the child, she may suffer social stress and, as a result of isolation from her own family and friends, have relatively restricted social interaction.

Taking care of a child is not an easy task, since it requires the execution of a complex, delicate and selfless tasks, which can impact the QL [20]. In this study, it was observed that a large number of the mothers in the G1 group had a family income between one and two minimum wages and had declared financial assistance from the government, such as the Bolsa Família and the social benefits due to the disability of the child.

In the context of QL, it is known that a stable socioeconomic condition is related to better QL, since the presence of financial problems may cause additional concerns [9]. QL can be affected by the difficulties found by residents from the state countryside, including sanitation and transportation, which were mentioned during the interviews.

Previous studies showed that the health conditions from the residents of the countryside are more precarious when compared to residents in the capital city. There is not always a water supply in the countryside. According to another study, even though half of the population have piped water in one of their rooms, only 18% of this water comes from the general distribution system. The remaining residences collect water from water wells, headwaters, and other sources. For human waste and residence toilets, waste disposal often occurs in an inadequate way, such as dumping of waste in rudimentary pits or direct ditches into rivers, lakes or the sea, which influences the health of the residents [15, 16].

In Brazil, the severely increased incidence of microcephaly is warning call to the urgent need of large investments aimed at improving the living conditions of the population. The lack of readily available water, a reality in the life of the residents from the countryside, predisposes water to storage for later use without proper care, creating favorable places for mosquito reproduction.

The study shows that in rainy periods, water can be dammed up in poor residences or where there is stored waste, creating favorable environments for ZIKV carrier proliferation. Stored garbage also becomes a perfect place for such proliferation. These factors show how the ZIKV outbreak may be related to the precarious living conditions of the population, and therefore, may be closely associated with the environmental and social determinants of health [21].

The mothers of the G1 group had lower QL scores. However, there was a statistically significant difference only for the environmental domain, which refers to physical security and protection, home environment, financial resources, health and social care, including availability and quality, opportunity to acquire new information and skills, participation and recreation/leisure opportunities, and the physical environment including pollution, noise, traffic, weather and transportation.

Another study with parents of hearing-impaired children, using the same questionnaire tool, also verified that the environmental domain of WHOQOL-bref was the most compromised [6]. This aspect is related to the housing and health conditions of the G1 group, especially the availability of transportation to take the child to the appointments. Although these differences were not statistically significant, it leads us to reflect on the lower QL scores of the G1 group, since the children had similar ages, the difference may not be directly related to the care of children, but rather to the complexity of care.

Children with eating disorders related to lactose, the G2 group in this study, can have gastrointestinal or immunological symptoms, which can lead to electrolyte imbalances or even trigger allergies. However, when treated, these changes can be tolerated and modified with the development of the individual [22, 23, 24]. This condition requires more attention from the mothers, however, in contrast to the G1 group, treatments are available which may allow symptoms to evolve over the course of these children’s lives.

With regard to these aspects, we can relate these characteristics to overburden of the mother in the care of other children in addition to the child with microcephaly, who needs more specific care.

Frustration with lack of sleep and leisure, due to a shortage of time and financial resources, are factors that impact the QL of these mothers. Some isolate themselves from society due to their personal appearance, which is a consequence of their lack of time due to being overburdened with the care with of their child, or because the child’s malformation actually changes their physical looks. This isolation will reduce the support these mothers can receive from society. In some cases, according to Almeida and Berlim [25, 26], isolation from society and being overburdened with the care of their child can lead to depression, which in turn impacts overall QL.

There was a statistically significant negative correlation regarding the number of children, pregnancies and births, related to all domains. When correlating maternal age, only the environmental domain and the total score were significant. Factors related to the environmental domain, such as difficulty in accessing services of basic needs (health, transportation, leisure, financial resources and safety), contribute to the increased level of stress of the caretakers. With regard to this aspect it was observed that the higher the maternal age, the lower the environmental score, therefore, maternal maturity and perceptions acquired with age may be related to each other.

We conclude that these mothers need greater attention, since they are overburdened with the care of their children, abdicating their own healthcare. The results obtained in this study indicate the need to develop programs for psychological support and education, and also improve living conditions, aiming to prevent any harm to the QL of these mothers.

The QL of the mothers of children with microcephaly had lower scores than the mothers of children with normal NPM, although the only significant difference was observed in the environmental domain. Environmental conditions were related to the housing and financial resources, and not necessarily to the children, which are required for the successful treatment and neuropsychomotor development of children. We emphasize that children should have routine monitoring and treatments with various specialists, and note that most of the subjects in this study relied on public transportation to receive these services.

A major limitation of this study was the number of participants, because it was not possible to recruit all the mothers who received these services due to their absence on appointment days and lack of specific laboratory tests, which associate microcephaly with ZIKV infection.

This study was of great importance, because it showed that the perception of QL of the mothers was not related to the NPM of the children. Still, it is of great relevance, since there are no studies specifically aimed at mothers of infants with this malformation.

We suggest new studies to monitor the mothers, since the future development of these children is still unknown. We believe that the burden of care and anxiety may increase as the children age. A longitudinal study would be ideal, since some mothers are still not aware that their children will not have a normal NPM.

The growth and development of these infants are influenced by various treatments, and fulfilment of routine care required for development will be the task of the maternal figure. We also suggest for future studies that the burdens of these caretakers be further evaluated.

